# Resolving a Guanine-Quadruplex Structure in the SARS-CoV-2 Genome through Circular Dichroism and Multiscale Molecular Modeling

**DOI:** 10.1101/2023.04.13.536758

**Authors:** Luisa D’Anna, Tom Miclot, Emmanuelle Bignon, Ugo Perricone, Giampaolo Barone, Antonio Monari, Alessio Terenzi

## Abstract

The genome of SARS-CoV-2 coronavirus is made up of a single-stranded RNA fragment that can assume a specific secondary structure, whose stability can influence the virus ability to reproduce. Recent studies have identified putative guanine quadruplex sequences in SARS-CoV-2 genome fragments that are involved in coding for both structural and non-structural proteins. In this contribution, we focus on a specific G-rich sequence referred as RG-2, which codes for the non-structural protein 10 (Nsp10) and assumes a guanine-quadruplex (G4) arrangement. We provide the secondary structure of the RG-2 G4 at atomistic resolution by molecular modeling and simulation, validated by the superposition of experimental and calculated electronic circular dichroism spectrum. Through both experimental and simulation approaches, we have demonstrated that pyridostatin (PDS), a widely recognized G4 binder, can bind to and stabilize RG-2 G4 more strongly than RG-1, another G4 forming sequence that was previously proposed as a potential target for antiviral drug candidates. Overall, this study highlights RG-2 as a valuable target to inhibit the translation and replication of SARS-CoV-2 paving the way towards original therapeutic approaches against emerging RNA viruses.

## Introduction

The outbreak and evolution of the COVID-19 pandemics, ^1–4^ caused by SARS-CoV-2 virus, has dramatically evidenced the risks related to emerging infectious diseases and the necessity to understand, at a molecular level, the complex mechanisms behind viral replication and reproduction. RNA viruses, including coronaviruses, can be counted among the most problematic emerging infectious diseases, due to their adaptability, high mutation rate, and transmissibility. A deep understanding of RNA viruses functioning, which in turn allows the rapid deployment of pharmaceuticals or vaccinal measures in case of an epidemic or pandemic event, is therefore highly needed.

SARS-CoV-2 viral genome is composed of a relatively large positive-sensed RNA strand of about 300k nucleobases. Upon cellular infection, and the entry into the host compartments, the genome is directly translated by cellular ribosome to produce the viral proteins which will further allow the reproduction of the virus and the diffusion of the infection. The sequencing and characterization of SARS-CoV-2 genome revealed that it codes for both structural proteins, including the well-known Spike (S),^5,6^ and non-structural proteins (Nsp) exerting enzymatic activity and favouring the viral maturation or its protection via the evasion of the host immune system.^7–10^ Among the non-structural proteins there are the viral proteases,^11,12^ which participate to the maturation of the originally produced large polyprotein, and the RNA-dependent RNA polymerase which, duplicating the viral RNA, allows its replication.^9,13,14^ Both proteins represent key targets for antiviral agents and have been largely studied by molecular and structural biology as well as simulation approaches.^13–15^

Despite its apparent simplicity, the genomic organization of RNA viruses hinders some level of complexity, often correlated with the virus reproductive capacity and infectivity.^16–19^ RNA and DNA viruses, including flaviviruses,^20^ Zika,^21^ Dengue,^22^ and human immunodeficiency viruses (HIV),^23^ possess putative guanine quadruplex sequences (PQS).^24,25^ These G-rich sequences are associated to crucial regulatory roles, especially in the framework of gene expression. Furthermore, the interplay between guanine quadruplexes (G4s) of the host cells and the viral protein machinery has important consequences on the virus life cycle. For instance, specific viral Nsp are activated by the interaction with G4s.^26,27^ The stabilization of such motifs with specific ligands hampers the replication or translation of the viral genome, blocking or slowing the viral infection.^28,29^ Thus, this approach represents an interesting and alternative therapeutic strategy to counteract emerging infectious diseases.^20,30^

G4s are arrangements forming in guanine-rich regions of RNA and DNA.^24,31^ They consist of π-stacked tetrads of four guanines locked by Hoogsteen-type hydrogen bonds.^32^ The guanines composing the tetrad do not need to be contiguous in the sequence and may be bridged by rather large and flexible loops. G4s are further stabilized by monovalent cations, usually Na^+^ or K^+^, which occupy the central channel generated by the stacked tetrads.^33,34^ In addition, G4s exist under different topologies, parallel, hybrid and antiparallel, depending on the relative orientation of the connecting strands and of the glycosidic bonds. In RNA, the ribose 2’-hydroxyl groups impose steric constraints that favour *anti* glycosidic bond angles. In turn, these facilitate formation of stable parallel topologies with shorter loops.^31,35,36^ Nevertheless, several antiparallel^37–39^ and hybrid^40,41^ RNA G4s have recently been reported indicating that these motifs present structural complexity not only within DNA. Parallel, antiparallel and hybrid G4 topologies are characterized by specific signatures in their electronic circular dichroism (ECD) spectra, thus allowing their recognition and characterization.^42–46^

Some PQS sequences have been recently spotlighted in SARS-CoV-2 genome.^29,47–49^ Qu et al.,^29^ focused their studies on one particular sequence, named RG-1, which codes for the nucleocapsid (N) protein, and is located in the region 28,903-28,917 of the viral genome. The stable G4 arrangement of RG-1 has been confirmed by ECD measurements^29,47^ and its precise atomic-scale structure has been predicted by our group using a multiscale simulation approach combining classical molecular dynamics (MD) simulations and hybrid quantum mechanics/molecular mechanics (QM/MM) techniques.^50^ Very recently, Qu et al. published a study on another PQS located in the region 3,467-3,483, forming a G4 which is more stable than RG-1 and suggested that this could be the target of a known G4 binder (TMPyP4) able to suppress SARS-CoV-2 infection in an infected animal model.^29^

A further PQS, labelled as RG-2, was identified in the region 13,385-13,404 and it codes for the Nsp10 of SARS-CoV-2.^29,47–49^ However, Qu et al,^29^ reasoning that RG-2 quadruplex is less stable than RG-1 and other G4s of the SARS-CoV-2 genome, did not consider it as valuable target for antiviral agents and they set it aside. On the contrary, we strongly believe that RG-2 could represent a more valuable target for G4-aimed drugs since it is more prone to stabilization. Furthermore, Nsp10 has a crucial role in assuring the viral maturation and represents a fundamental cofactor in the activation of multiple replicative enzymes.^51^ Indeed, the formation of a complex with Nsp10 is necessary to assure the exoribonuclease activity of Nsp14, which constitutes a proof-reading step avoiding RNA replication errors.^52^ Nsp10 is also necessary to guarantee the methyltransferase capacity of Nsp16, which increases the RNA stability by its capping with a methyl group.^53^ The indirect inhibition of the replication and repair-capacity of Nsp14 by targeting the RG-2 G4 may also increase the efficiency of nucleoside-analogous antiviral agents.

Globally, the possibility to develop G4-targeted antiviral agents against emerging infectious diseases, and RNA viruses in particular, is highly appealing.^47^ However, to allow a rational design of potential therapeutic agents, hence to maximize their efficiency and selectivity, it is necessary to obtain a proper atomistic-resolved structure of the involved targets. In this respect, the determination of RNA secondary structures may be challenging also due to the inherent instability of RNA oligomers. In a previous contribution^50^ we have reported a multiscale approach which has confirmed the G4 arrangement of the SARS-CoV-2 RG-1 sequence, providing its full-atom structure. This was achieved by long-range MD simulations on a starting guess arrangement obtained by sequence alignment between RG-1 and other G4 sequences. The simulation of the ECD spectrum on top of snapshots issued by the MD simulation has provided a one-to-one mapping with the experimental results, confirming the soundness of the proposed model-structure.^50^

## Results

In the present contribution we provide the first structural resolution prediction of the RG-2 G4 in SARS-CoV-2, relying on ECD spectroscopy and exploiting a similar computational protocol to describe its arrangement at the atomic-level. The simulated ECD spectra, obtained at hybrid quantum mechanics/ molecular mechanics (QM/MM) level is also used to support the soundness and robustness of the computational model. In addition, the stabilization of the G4 arrangement of RG-2 also in comparison with RG-1, by pyridostatin (PDS), a well-known G4-ligand,^54^ is assessed by FRET melting analysis and ECD. The full experimental and computational protocols are reported and discussed in the Electronic Supplementary Information (ESI).

The RG-2 segment of the SARS-CoV-2 genome is a guanine rich area in the Nsp10 coding-region and is characterized by the 3’-GGUAUGUGGAAAGGUUAUGG-5’ primary sequence.^29,47–49^ The alignment between RG-2 and different G4-forming sequences has been performed and the maximum similarity has been obtained with the PDB codes 3CDM^55^ and 6E8S^40^, a parallel DNA and hybrid RNA G4, respectively (see ESI and Table S1-S2 and Figure S1-S2). Considering that experimental results recently published ^29,47–49^ indicate that RG-2 adopts a parallel folding, the structure corresponding to the PDB code 3CDM has been chosen as a starting template, after converting DNA to RNA. Subsequently, two independent MD replicas, each exceeding the μs-time scale, have been performed, using the NAMD code.^56,57^

After a slight structural reorganization observed in the very first steps of the simulation, RG-2 evolves towards a stable parallel G4 arrangements composed of two stacked tetrads and stabilized by a central K^+^ ion (Figure 1).

**Figure 1.**
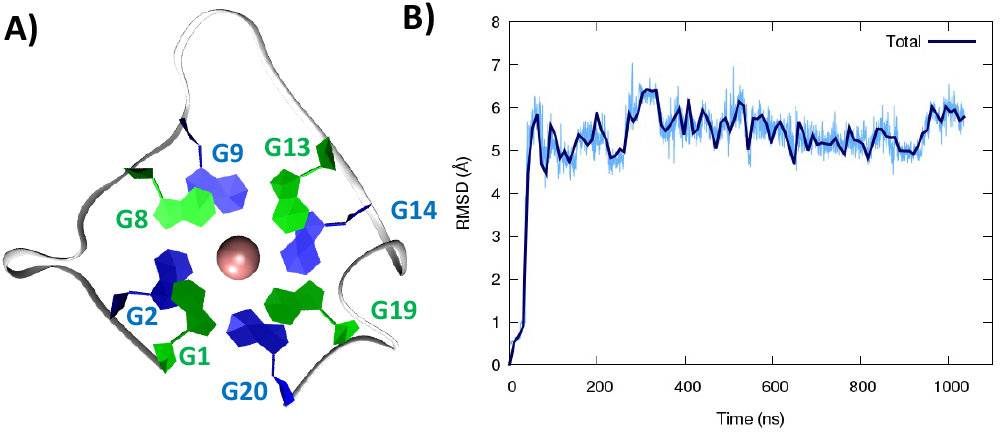
A representative snapshot of the parallel G4 formed by the RG-2 sequence as obtained by MD simulations. Guanine residues composing the two superposing tetrads are evidenced by different colors (A). The corresponding time series of the RMSD for the RNA strand (B). Analogous results obtained for the second replica are reported in Figure S3A.

As seen in Figure 1A, the first tetrad is composed by G1, G8, G13, and G19. The residues G2, G9, G14, and G20 constitute the second tetrad. The stability and the rigidity of the structure can be also appreciated by the time evolution of the Root Mean Square Deviation (RMSD), which is reported in Figure 1B. After the initial increase the RMSD remains stable throughout the whole simulation. The two tetrads are bridged by rather short, yet still flexible unstructured loop. Interestingly, this aspect constitutes a significant difference with the RG-1 sequence, which instead presents a long and extremely flexible loop responsible for the global G4 conformational flexibility.^50^ The absence of long peripheral loops is enhancing the rigidity of the RG-2 arrangement, while providing a globally more compact structure as can be seen in the superposition of the snapshots extracted from the MD trajectory shown in Figure 2A. Unsurprisingly, the G4 arrangement also leads to a rather rigid core, whose structural parameters are close to the ideal geometry. The two G4 tetrads experience a twist angle between their axis close to ideality averaging at 30.8±2° for the first replica (Figure 2B) and at 29.0±2° for the second one. The same rigidity can also be appreciated by the distance between the centre of mass of the two tetrads, which also shows a very limited evolution and oscillation all along the simulation comprised between 3.43±0.09 and 3.45±0.08 Å, for the first and the second replica, respectively (Figure 2C). These values are also coherent with near ideal π-stacking between the tetrads, globally confirming the overall stability of the aggregate. Finally, as can be appreciated in Figure 2D, the twist angle between the main axis of the tetrad-forming guanine residues is also rather rigid, as exemplified by the limited spread of the distribution.

**Figure 2.**
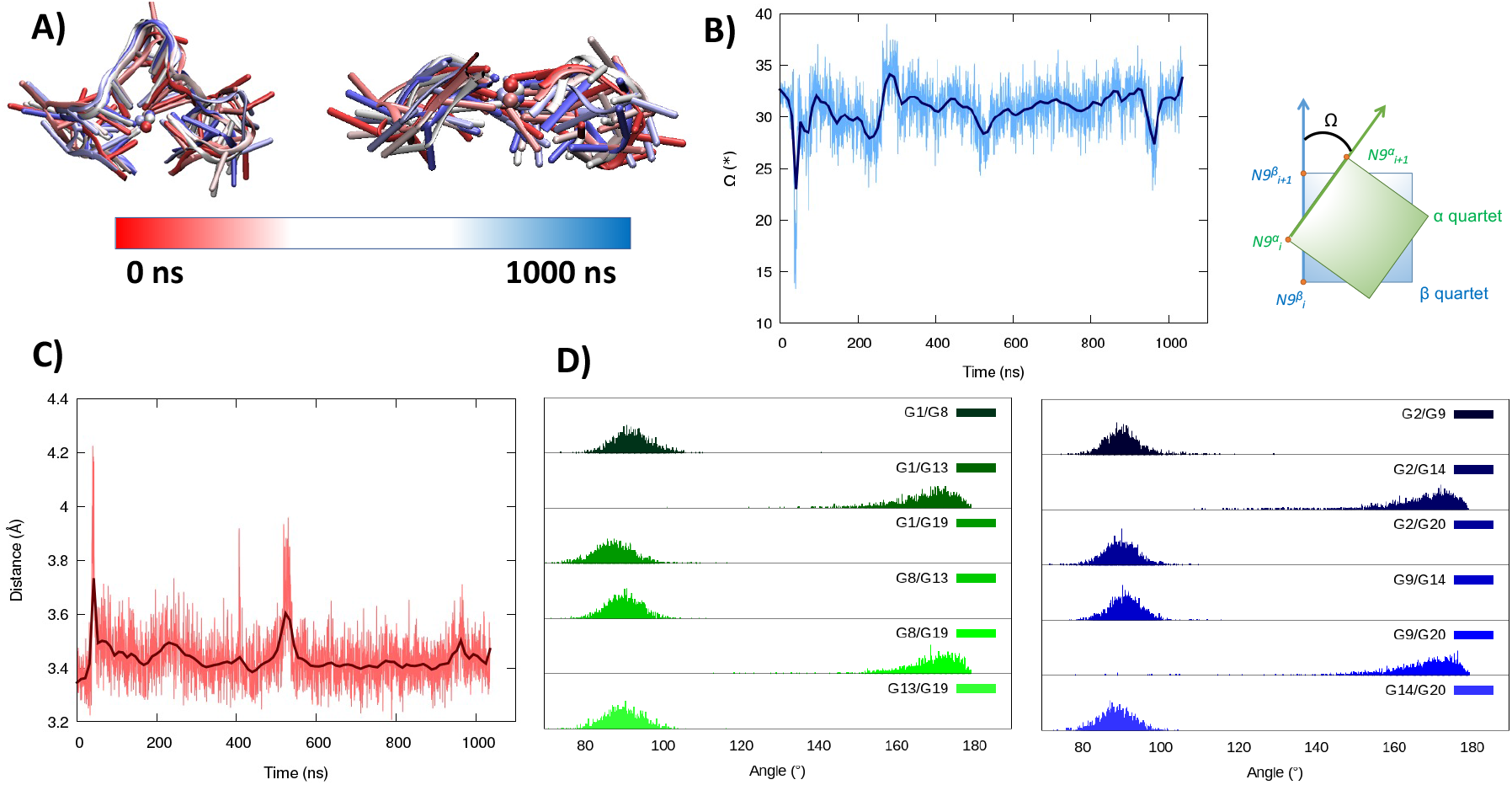
Evolution of the structure of RG-2 in its parallel arrangement along the MD simulation in top and front view (A). Time series of the twist angle Ω between the tetrad and its geometrical definition (B). Time series of the distance between the tetrads (C). Distribution of the angles between the axis of the guanines composing the first, green, and the second, blue, tetrad (D). Analogous results obtained for the second replica are reported in ESI (Figure S4). For the precise definition of the involved axis see the original work by Tsetkov et.al.^58^

Considering the alignment results described in SI, we built a model based on a hybrid G4 structure (PDB code 6E8S)^40^and performed MD simulations. As for the parallel arrangement, following a minor structural reorganization noted during the initial stages of the simulation, RG-2 forms a very stable hybrid G4 consisting of two stacked tetrads stabilized by the presence of a central K^+^ ion (Figure 3).

**Figure 3.**
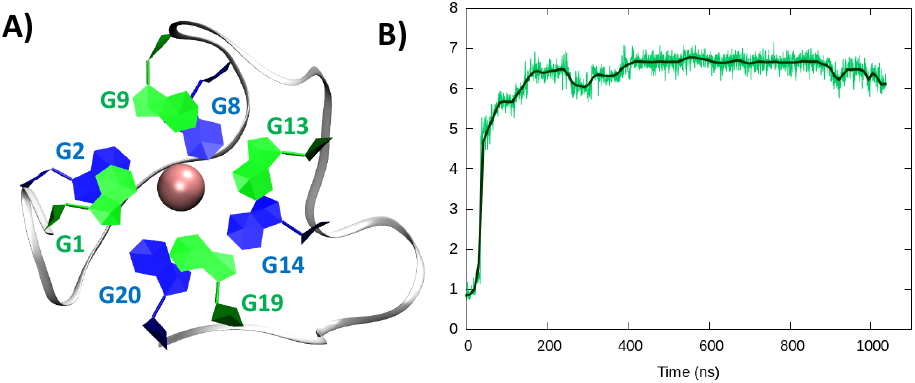
A representative snapshot of the hybrid G4 formed by the RG-2 sequence as obtained by MD simulations. Guanine residues composing the two superposing tetrads are evidenced by different colours (A). The corresponding time series of the RMSD for the RNA strand (B). Analogous results obtained for the second replica are reported in in Figure S3B

In the hybrid structure, the first tetrad is composed by G1, G9, G13, and G19, while the second by residues G2, G8, G14, and G20. All the considerations made for the parallel arrangement hold true also for the hybrid folding. The time evolution of the RMSD (Figure 3B) reveals the formation of a stable structure with slightly less oscillation than the parallel. Looking at the structural parameters (Figure 4), the simulation yields an ideal G4 arrangement. The twist angle of the two G4 tetrads averages at 35.1±3° for the first replica (Figure 4B) and at 27.5±4° for the second one. Also, the distance between the centre of mass of the two tetrads results to be 4.00±0.5 Å, with slightly less oscillation compared to the parallel structure (Figure 4C). All along the simulation, the spread of distribution of the angles between the guanines is quite limited (Figure 4D).

**Figure 4.**
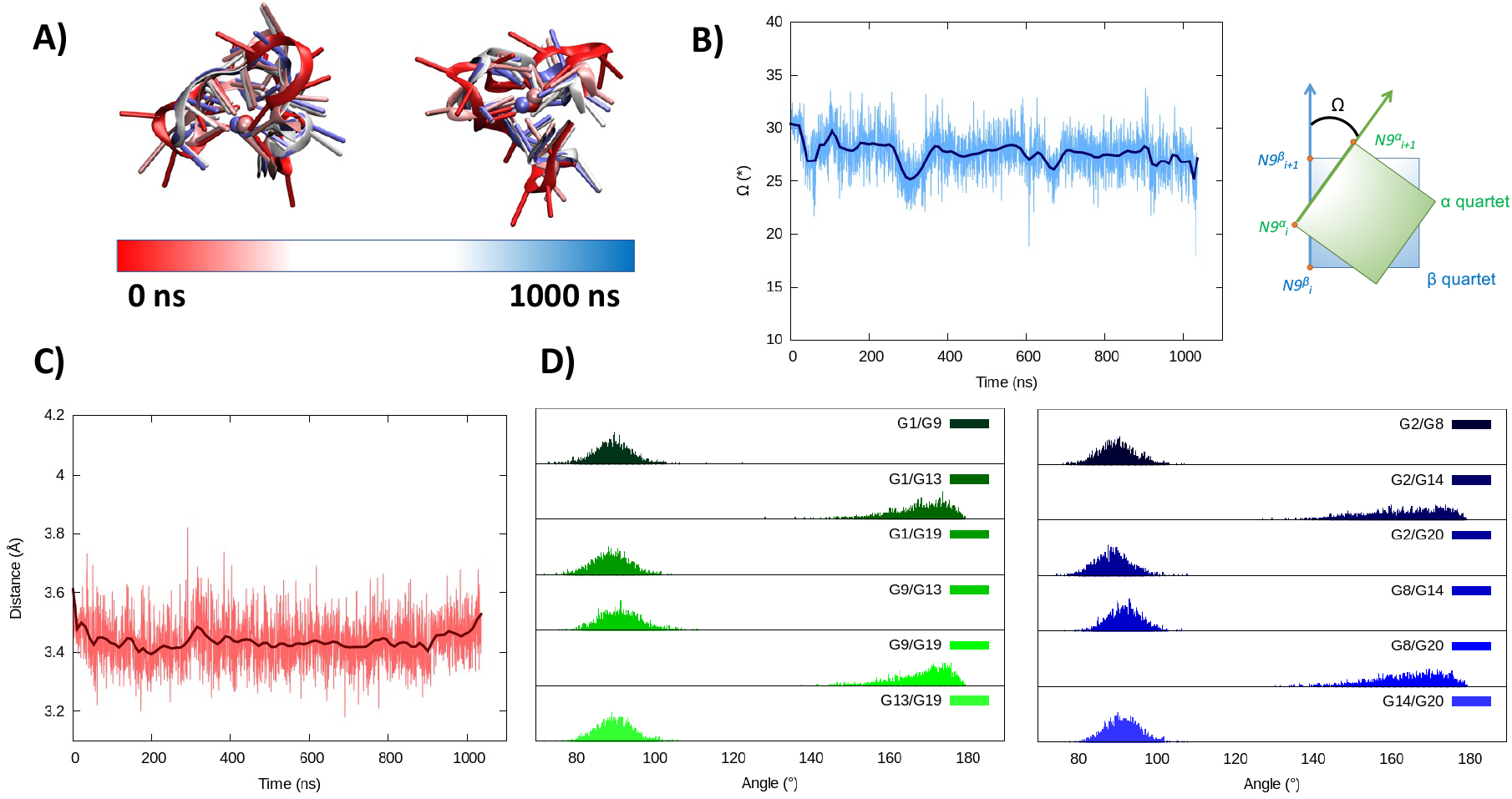
Evolution of the structure of RG-2 in its hybrid arrangement along the MD simulation in top and front view (A). Time series of the twist angle Ω between the tetrad and its geometrical definition (B). Time series of the distance between the tetrads (C). Distribution of the angles between the axis of the guanines composing the first, green, and the second, blue, tetrad (D). Analogous results obtained for the second replica are reported in ESI (Figure S5)

Taking into account that both structures, parallel and hybrid, coming out of the simulation are very stable, to confirm the G4-arrangement of RG-2 we recorded the ECD spectrum of the corresponding G-rich sequence folded in K^+^containing solution and compared to the simulated ones (Figure 5 and S6-S7).

**Figure 5.**
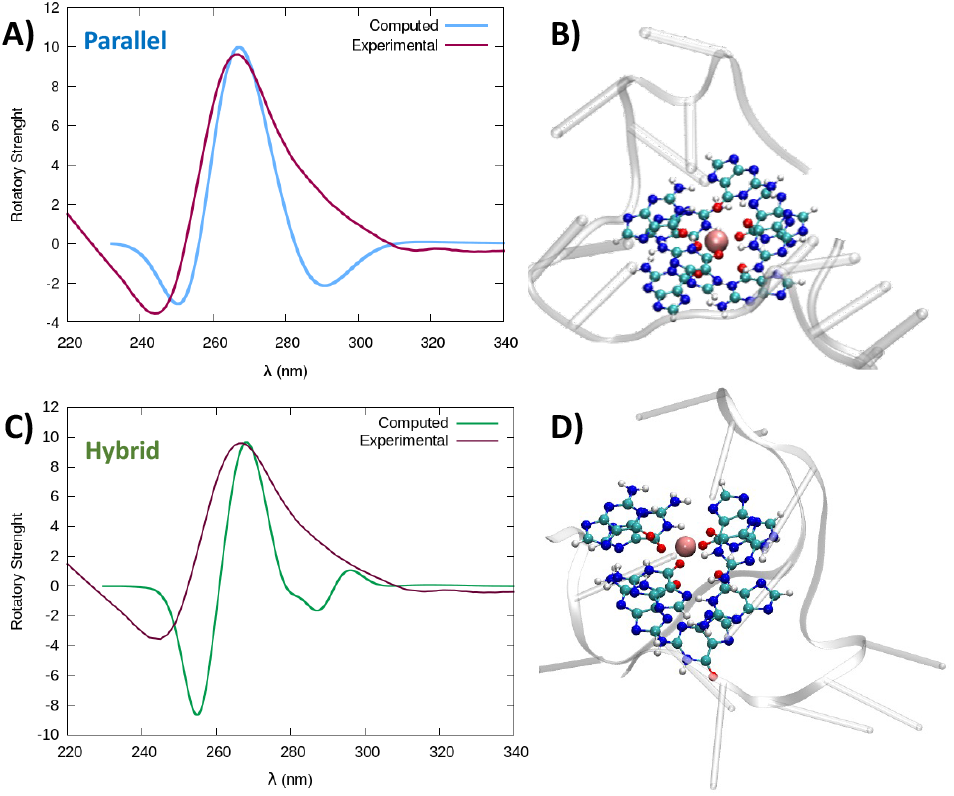
Comparison between the experimental and simulated QM/MM ECD spectrum of parallel and hybrid RG-2 structures in water solution (A and C, respectively). The computed spectrum is simulated by QM/MM calculations, using the M06-2X functional, similar results are obtained using the ωB97XD, see ESI. To reproduce the experimental signal a broadening factor of 0.15 eV has been applied to each vertical transition. Note that vertical transition in the context of Born Oppenheimer approximation represent transition to the electronic excited states without any geometrical change of the nuclei. The QM partition is illustrated in panels B and D in ball and stick representations and comprises all the nucleobases forming the tetrads and the central K^+^ ion.

The experimental spectrum exhibits a weak negative band at 240 nm, a strong positive ellipticity at 260 nm followed by a positive shoulder at 290 nm. Our result is more in line with the one published by Zhang et al.,^49^ while Qu and collaborators, in both of their papers,^29,47^ do not report about the presence of the little shoulder at around 290 nm.

The G4 topology was identified by using the conformation index “r” introduced by Jean-Louis Mergny and colleagues (see experimental section in ESI).^59,60^ In our experimental conditions, the “r” value for RG-2 is significantly higher than 0.5 (0.7), providing evidence for a predominant parallel conformation. To support this finding, we employed the software developed by John O. Trent, Jonathan B. Chaires and collaborators, which utilizes an algorithm to quantitatively estimate the secondary structure content of quadruplexes based on their experimental CD spectra.^46^ The results obtained by fitting our experimental data with this software confirm the predominant presence of a parallel arrangement in RG-2 (see ESI and Figure S8).

The simulated spectrum, which is obtained considering the evolution of the sequence geometry along the MD simulation for the parallel arrangement, matches satisfactorily with the shape of the experimental ECD. However, the shoulder appearing at around 290 nm is missing, coherently with the fact that our model only involves a parallel arrangement, while the ensuing negative signal at around 240 nm is slightly underestimated in our simulation.

On the contrary, the slight shoulder at 290 nm is present in the simulated spectrum of the hybrid structure, which surprisingly retains the intense band at around 260 nm. It is generally accepted that parallel-type G4s show a positive ECD band at around 260 nm, due to the *anti*-*anti* guanines that stack over each other. A strong peak centred at around 295 nm, due to *syn*-*anti* guanines, is typically associated to antiparallel structures. It must be said that there are different cases for which the use of those ECD signatures to distinguish parallel from antiparallel or hybrid folding topologies does not always lead to the expected result.^61–66^ For instance, hybrid structures where three strands are stacking *anti*-*anti* and one strand is stacking *syn*-*syn*, result in ECD spectra characterized by a positive peak at 260 nm (with a slight shoulder at 290 nm) and negative band at 240 nm, which generally indicate predominant parallel G4s.^61^ As it can be verified by processing the representative snapshots structures (available in ESI) in webservers like ASC-G4 or WebTetrado,^67,68^ our optimized hybrid G4 structure is characterized by the following configuration: G1-G2 *anti-anti*; G9-G8 *syn-syn*; G13-G14 *anti-anti*; G19-G20 *anti-anti*. This is exactly the case when hybrid structures give ECD spectra similar to the one experimentally recorded. The hybrid RNA G4 with PDB code 6E8S has also three strands where guanines are stacking *anti-anti* and one strand is stacking *syn-syn*, although its ECD was not recorded since the G4 is part of a bigger structure where a duplex is also involved. Additionally, the hybrid structure is characterized by two lateral loops that can also be associated to the presence of the shoulder at 290 nm.^46^

Globally, our results confirm that RG-2 sequence can equally form stable parallel and hybrid G4 structures. The analysis of the structural parameters and the comparison of calculated and simulated ECD spectra, as well as important similarities with the 2-quartet G4-RNA with PDB code 6E8S, indicate a slight preference for the hybrid structure. We note that a shift of the absorption wavelengths has been applied to the calculated spectrum. As already discussed in our previous contribution,^50^ the blue-shifted computed ECD signal is mainly due to incomplete basis set but does not prevent comparing the shape of the main transition to validate the structure topology assignation.

We also recorded the ECD spectrum of RG-2 sequence in Na^+^ containing buffer and in the presence of PEG-200 as crowding agent. The spectrum in in Na^+^ ions is characterized by the presence of a strong positive band at 260 nm, (Figure 6), while the shoulder at 290 nm is no longer observed. According to what discussed above, this could indicate the predominance of the parallel structure. Interestingly, the ECD recorded in K^+^ buffer with 40% PEG-200 shows a noticeable shift for the positive band at 258 nm and the presence of a weaker band at around 300 nm. These features are typical of an antiparallel topology of group III, according to the classification proposed by Webba da Silva and collaborators.^42^

**Figure 6.**
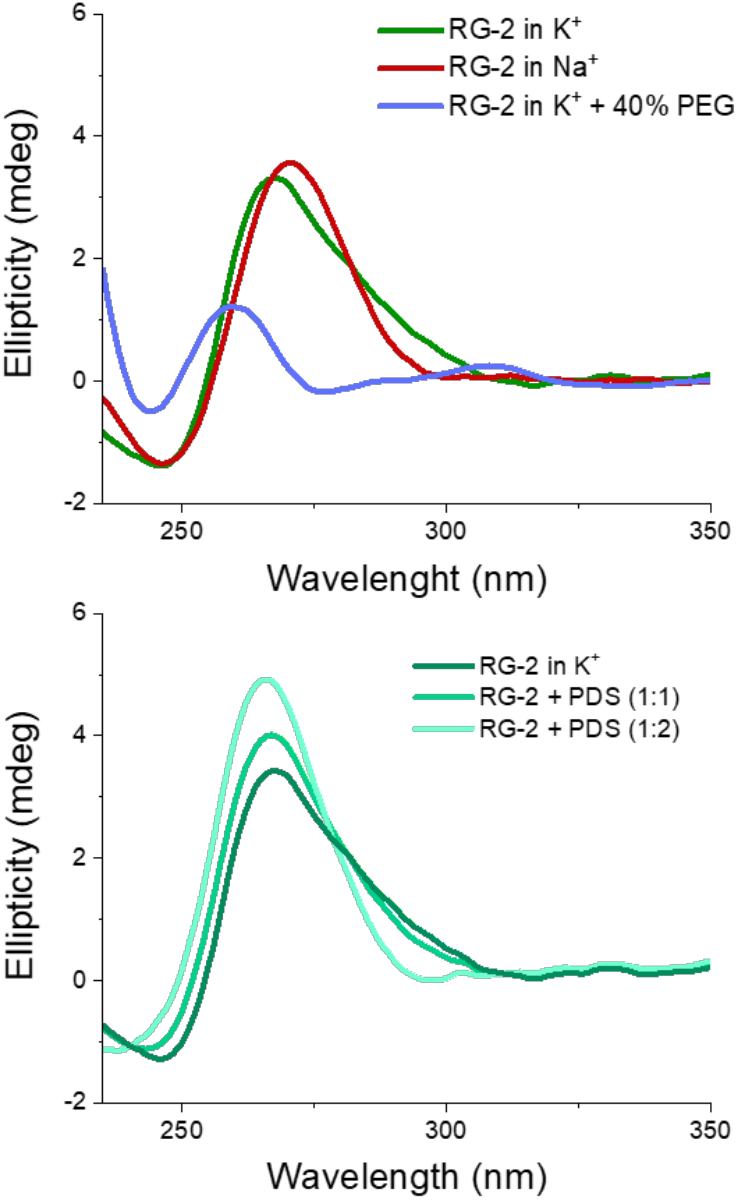
Upper panel: experimental ECD spectra of RG-2 in K^+^ or Na^+^ containing buffers and in the presence of PEG-200 as crowding agent. Lower panel: experimental ECD spectrum of RG-2 in K+ in the presence of increasing concentration of PDS.

The spectra of RG-2 were subsequently recorded in various buffers while in the presence of PDS, a well-known binder of G4, in order to detect any potential alterations resulting from their interaction. Two different [RNA]:[PDS] ratios were used (1:1 and 1:2, respectively). As reported in Figure 6 (lower panel), when PDS is added to RG-2 in K^+^ buffer the intensity of the ECD band increases according to the concentration of the ligand followed by the disappearance of the shoulder at 290 nm. This result indicates the binding of PDS to RG-2. The alteration observed in the CD spectrum could potentially arise from the induced CD bands of PDS itself, as it absorbs in a similar range as the quadruplex, including within the 280-300 nm wavelength range. However, in Na^+^ buffer and in the presence of the crowding agent PEG, the increasing concentration of PDS produces only slight changes of the G4 spectrum (Figure S9-S10).

The formation of a stable aggregate between RG-2 and PDS has also been confirmed by MD simulations as shown in Figure 7. To this aim we simulated the self-recognition between both parallel and hybrid G4 conformations of RG2 and PDS by placing the latter in the water bulk and letting the ligand and the nucleic acid evolve spontaneously, without any external bias. Indeed, after equilibration, PDS is rapidly attracted toward the G4 and forms a highly stable complex via π-stacking with the exposed tetrads. Interestingly, the time evolution of the distance between the centre of mass of PDS and of the tetrad-forming guanines remains remarkably stable for a time-laps exceeding the μs scale in both simulations, with enhanced stability for the hybrid-PDS complex which shows slightly less oscillations. The structural parameters of the G4-folded RG2 are only slightly perturbed by the presence of PDS (see ESI Figure S11 and S12), justifying the experimentally observed similarity between the ECD spectra.

**Figure 7.**
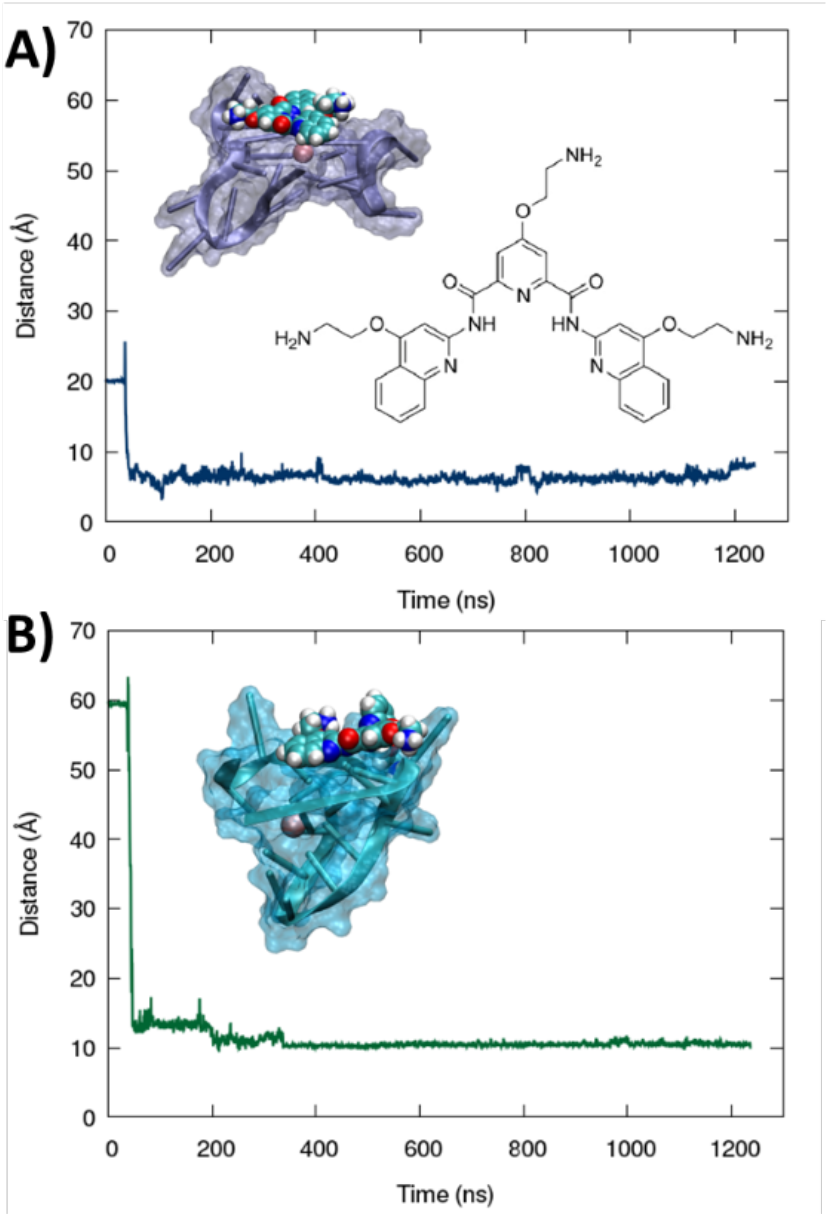
Time evolution of the distance between the centres of mass of PDS and the guanine-forming tetrads of the RG-2 sequence along MD simulation for both parallel (A) and hybrid (B) structures. The sharp peak at short time corresponds to the initial non-interacting conformation. A representative snapshot is also provided in the inlay, the G4 is represented in cartoon and transparent surface representation while PDS is displayed in van der Waals style. The chemical structure of PDS is also given in the figure.

As mentioned before, Qu et al.,^47^ comparing RG-2 and RG-1 G4 melting temperatures measured by ECD, concluded that RG-2 is less stable under physiological conditions showing a T_m_ around 45 °C. For this reason, the authors focused their studies on RG-1 only, deeming this G4 a better target for antiviral drug candidates. We performed Forster Resonance Energy Transfer (FRET) melting assay using RG-1 and RG-2 oligonucleotides, modified with a fluorophore (FAM) and a quencher (TAMRA) at the 5’ and 3’ ends, respectively, alone or in presence of increasing concentration of PDS. Contrary to what reported by Qu et al.,^29^ in our experiment RG-1 and RG-2 have almost the same melting temperature 51.4 and 48.7 °C, respectively, with RG-2 slightly less stable but certainly compatible with G4 folding in physiological conditions. The difference with data from Qu et al.^29^ can be attributed to the slightly different experimental conditions (type of buffer, K^+^ concentration and presence of fluorophores). However, and interestingly, the RG-2 quadruplex is stabilized more efficiently by PDS when compared to RG-1 at 1:5 [RNA]:[PDS] ratio (ΔT_0.5_ = 9.5 and 2.6, respectively, Figure 8). Therefore, its targeting for antiviral purposes appears highly suitable.

**Figure 8.**
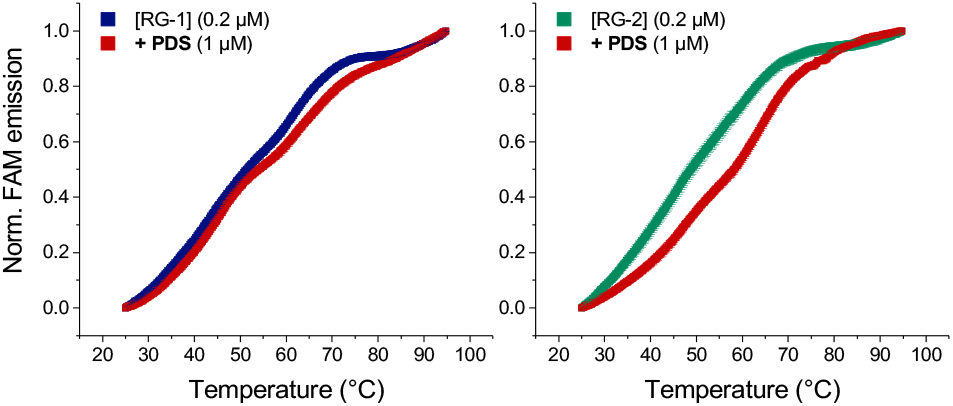
FRET melting curves of RG-1 and RG-2 (potassium cacodylate buffer, pH = 7.4) in the presence of PD

## Conclusions

In this contribution we present the first structural prediction of a SARS-CoV-2 genome sequence involved in coding the Nsp10 protein, which is essential for viral replication. In particular, we have shown that RG-2 can assume both stable parallel and hybrid G4 arrangements, composed of two rigid tetrads, almost ideally π-stacked, bridged by short, yet flexible loops. The analysis of the structural parameters, the simulation of the ECD spectra and their comparisons with the experimental data indicate a slight preference for the hybrid structure.

We confirm that our protocol, based on the combination of multiscale modelling, including sequence homology, MD simulations, and in silico spectroscopy, combined with experimental ECD spectroscopy represents a very suitable solution for the proposition of secondary structures in the genome of emerging viruses. Thus, it will offer a robust strategy for the rational development of antivirals targeting emerging infectious diseases. Furthermore, starting from our proposed models, we are working on the experimental determination of the RG-2 quadruplex.

We have also shown by ECD and FRET that the well-known G4-binder PDS stabilizes RG-2 secondary structure more efficiently than RG-1, offering a valuable antiviral strategy against emerging RNA viruses. Indeed, the stabilization of RG-2 G4 would result in the inhibition of the RNA translation and replication, and hence will hamper the viral diffusion. This is particularly true since Nsp10, which RG-2 participates to encode, is involved in critical processes assuring SARS-CoV-2 fitness and is related to the proof-reading of the RNA synthesized by the RNA-dependent-RNA-polymerase, and to the increase of its stability by methyl capping. In perspective, we aim at evaluating how Nsp10 expression changes in SARS-CoV-2 infected host cells to further validate RG-2 as a suitable target for COVID-19 treatment. Considering that G4 forming sequences, including RG-2, are highly conserved among coronaviruses,^49^ our definitively paves the way for possible efficient treatment in the fight against new emerging viruses of this family.

## Author Contributions

Luisa D’Anna: conceptualization, investigation, methodology, formal analysis, visualization, writing - review & editing; Tom Miclot: investigation, methodology formal analysis, visualization, writing - review & editing; Emmanuelle Bignon: investigation, methodology, formal analysis, writing - review & editing; Ugo Perricone: investigation, formal analysis, writing - review & editing; Giampaolo Barone: conceptualization, writing – original draft, methodology, formal analysis, visualization; Antonio Monari: conceptualization; writing – original draft methodology, formal analysis, visualization; Alessio Terenzi: conceptualization, writing – original draft, methodology, formal analysis, visualization. @

### Conflicts of interest

There are no conflicts to declare.

## Acknowledgements

All the simulations have been performed on the LPCT, Explor, and GENCI computing resources which are gratefully acknowledge. Financial support from the French Ministry for Higher Education and Research (MESR) and CNRS is also acknowledged. A.M. thanks A.N.R. and CGI for their financial support of this work through Labex SEAM ANR 11 LABX 086, ANR 11 IDEX 05 02. The support of the IdEx “Université Paris 2019” ANR-18-IDEX-0001. This work was also financed by European Union – NextGenerationEU – fondi MUR D.M. 737/2021 – project PRJ-0989 (A.T.). A.T. also thanks FFR-D15-162636 project.

## Notes

### Competing Interest Statement

The authors have declared no competing interest.

### Summary of Updates

Taking into account hybrid G4 structures by molecular modeling.

