## Supplementary Information for "Resolving a Guanine-Quadruplex Structure in the SARS-CoV-2 Genome through Circular Dichroism and Multiscale Molecular Modeling"

a) Department of Biological, Chemical and Pharmaceutical Sciences, University of Palermo, 90126 Palermo, Italy;
b) Université de Lorraine and CNRS, UMR 7019 LPCT, F-54000 Nancy, France;
c) Fondazione Ri.MED, Via Filippo Marini 14, 90128, Palermo, Italy
d) Université Paris Cité and CNRS, ITODYS F-75006 Paris, France

Corresponding authors
G.B., A.M., A.T.

### Computational Methodology

*Search for homologous G-quadruplex structures in the RCSB PDB databank as template for the parallel RG2 arrangement*

RG-2 consists of 20 nucleotides that, according to recent findings, should adopt a parallel G4 structure.[1–3] Thus, we used the strategy adopted in a recent contribution.[4] Specifically, the RG-2 RNA sequence was converted to DNA and the structure of the human telomeric G4 (PDB:1KF1) was used as a template to find a parallel G4, enforcing the "Relaxed structure" and "PROSITE" parameters. The best results are reported in the Table S1.

**Table S1** DNA parallel G4 homologous to RG2 found after searching in the RCSB PDB database.

| PDB code | Organism | Name | Sequence |
|---|---|---|---|
| 1KF1 | Human | Telomeric DNA quadruplex | AGGGTTAGGGTTAGGGTTAGGG |
| 3CDM | Human | Telomeric DNA quadruplex | TAGGGTTAGGGTTAGGGTTAGGG |
| 2M27 | Human | VEGF promoter | CGGGGCGGGCCTTGGGCGGGGT |
| 6JWE | Human | oncogene RET G-quadruplex | GGGGCGGGGCG-GGGCGGGGT |
| 5W77 | Human | MYC G-quadruplex | TGAGGGTGGGTAGGGTGGGTAA |
| 6T51 | Human | KRAS22RT promoter | AGGGCGGTGTGGGAATAGGGAA |
| 6ZRM | Human | RANKL gene | TGGGAGGGAGCGGGAGTGGG |
| 2N4Y | HIV-1 | synthetic construct (32630) | CTGGGCGGGACTGGGGAGTGGT |

*Refinement of the parallel G4 provided by the homology search*

The best candidate structure to be used as a template for the reconstruction of the RG-2 model was chosen based on a sequence alignment performed using the server version of M-Coffee.[5] Each of the G4 sequences were aligned with the sequence of RG-2 converted to DNA (Figure S1-A). The sequences of PDB:3CDM and PDB:6ZRM and PDB:5W77, showing the best scores, have been kept. Besides sequence similarity, structural features were also considered as criterion for the selection. Therefore, each G4 was examined to locate the position of loops, tetrad-forming guanines, bulges, and nucleotides not participating in the formation of the G4 main structural elements. This was done by manually aligning the nucleotide sequences according to their structures (Figure S1-B). G4 presenting bulges were eliminated to avoid instabilities in the reconstruction process. Thus, PDB:6T51, PDB:6ZRM and PDB:2N4Y have been excluded from the panel of potential templates. After this step we considered the G4 whose loops size is similar to those of RG-2, which leaves only one potential candidate, PDB:3CDM, which was thus selected.

```
A) Sequence alignment
  RG-2|SARS-CoV-2|RNA converted to DNA        -GGT--ATGTGGAAAGGTTATG--G
  3CDM|Human|telomeric DNA quadruplex         TAGGGTTAGGGTTAGGGTTAGG--G
  2M27|Human|VEGF promoter                    CGGG--GCGGGCCTTGGGCGGGG-T
  6JWE|Human|oncogene RET G-quadruplex        -GGG--GCGGGGCG-GGGCGGGG-T
  5W77|Human|MYC G-quadruplex                 TGAGGG--TGGGTAGGGTGGGTA-A
  6T51|Human|KRAS22RT promoter                -AGG--GCGGTGTGGGAATAGGGAA
  6ZRM|Human|RANKL gene                       TGGG--AGGGAGCGGGAGTGG---G
  2N4Y|HIV-1|synthetic construct (32630)      CTGG--GCGGGACTGGGGAGTGG-T

B) Structure alignment
  RG-2|SARS-CoV-2|RNA converted to DNA        ---GG-TATGTGG-AAA-GG---TTATGG------
  3CDM|Human|telomeric DNA quadruplex         TA-GGGTTA--GGGTTA-GGG--TTA-GGG-----
  2M27|Human|VEGF promoter                    CG-GGGC----GGGCCTTGGG--C---GGG--GT-
  6JWE|Human|oncogene RET G-quadruplex        ---GGGGCG--GGGC---GGG--GCG-GGG--T--
  5W77|Human|MYC G-quadruplex                 TGAGGGT----GGGTA--GGG--T---GGG--TAA
  6T51|Human|KRAS22RT promoter                A--GGGC----GG-T---GTGGGAATAGGG--AA-
  6ZRM|Human|RANKL gene                       T--GGGA----GGGAGC-GG---GA--GTGGG---
  2N4Y|HIV-1|synthetic construct (32630)      CT-GGGC----GGGACTGGG--A---GTGG-T-
                                                    L1        L2        L3
```

**Figure S1** A) Sequence alignment obtained using M-Coffee and B) structure alignment highlighting the guanines forming tetrad (Green), the three loops L1-3 (Pink), bulges (Yellow) and the nucleotides not participating to the G4 folding located at the 5'- 3'-termini (Grey).

### *Search for structurally resolved RNA sequences in the RCSB PDB database*

The homology search strategy used previously did not provide satisfactory results when considering RNA structures. The same is true when the RG2 sequence is used in the advanced search of the RCSB PDB website (https://www.rcsb.org/). Consequently, the search method consisted in spanning all resolved RNA structures contained in the RCSB PDB. The G4s presenting the closest similarity to RG2 sequence have been selected. Notably, we restricted the search to RNA nucleic acids only, while DNA/RNA mixed structures have been excluded. The search returned 1820 results which were exported as FASTA files.

### *Search for RNA G4s displaying hybrid topology with the most similar sequence to RG2*

In this step, the RG2 RNA sequence is compared with all RNA sequences contained in the previously generated FASTA file. To do this, we used the BLAST method.[6] The *Align Sequences Nucleotide BLAST* online tools provided by NCBI[7] has been used with Megablast.[8] The top five hits are reported in Table S2:

**Table S2.** RNA hybrid G4 homologous to RG2.

| PDB code | Organism | Name | Sequence |
|---|---|---|---|
| 7PTQ | Synthetic | RNA origami 5-helix tile | GGAAUUAGAGUGUGUUCCUGAACUGCUUCGGCGGUUCG CUACGUUCUUCGGAAUGUAUAUAGUGUUCGCAUUAUAC CGUAGUCCAAGCCGUGUGGUUCGCCGCACGUCGUUCGUU CGCGGACGAGCAGGUGCCAUAACCUCCAAAUGGUACCUG CGCGUGUUGUCAGCAAGGUCUAAGCUGAUAACACUAUGC UAACGACUGAAGCAUAUUGGAUUACGGGCCAAGGGCAGC UAAGAUCGGAAGCUGUCCUUGAGGAACGCACUCUGAUUC CCCUCCGGAAGGAGCCCCACAGGUAAUAAACCGAUCAUAU UACUUGUGCACUCGCAACAGUCGAGCGGGUGGUAAUGAU UGCGCCCGUUGGCUAGAAUAGACCACUAGCUAACGGCGG GUCUUGGAUCAAUGGAGGAGAUCCAGGACCCGACCCGGA CUUCGGUUCGGGUGCGGCCUUAGUUCGCUGAGGCCCGUC GCAUUCGUGUGACGGUGGUUAUUGCGGUCAAGGUUUCG ACUUUGAGUAUUCCUUCGGGGAUACGCUUCUUCUGGAG G |
| 6UP0 | Synthetic | RNA Mango-III fluorescent aptamer | GCUACGAAGGAAGGAUUGGUAUGUGGUAUAUUCGUAGC |
| 6E8T | Synthetic | RNA Mango-III (A10U) aptamer | GUACGAAGGAAGGUUUGGUAUGUGGUAUAUUCGUAC |
| 6E8S | Synthetic | RNA iMango-III aptamer | GCUACGAAGGAAGGAUUGGUAUGUGGUAUAUUCGUAGC |
| 7SHX | Human | a non-coding RNA transcript | GGACCCAUAACCCACCUAGACCCUAGCUUCGGCUAGAGGG UCAACGCGAAAGCGAGGCCGGGUGGGCGGGUACGUCCGC CCUGGGGAGGGGUCC |

Since the only G4 structures of interest are 6UP0, 6E8T, 6E8S, only these have been retained (Figure S2).

```
RG-2   1-GGUAUGUGG-9
6UP0 18-GGUAUGUGG-26   Score 18.3 Expected 4.5
6E8T 17-GGUAUGUGG-25   Score 18.3 Expected 4.5
6E8S 18-GGUAUGUGG-26   Score 18.3 Expected 4.5
```

**Figure S2**. Blast sequence alignment between RG2 and the 3 top hits RNA G4

#### *Selection of the hybrid G4 structure used as template for RG2*

The BLAST alignment results show that the selected three structures align exactly on the same part of the RG2 sequence and return the same score. Since sequence alignment is unable to distinguish between them, it is necessary to consider structural features. However, their structure is also highly similar consisting of a G-quadruplex-coiled strand of RNA overlaid by a helical portion as also confirmed by the DSSR-G4DB database.[9] Structure 6E8T has been discarded because the nucleotide 11 is absent from the resolved structure available in RCSB PDB. Since the two remaining structures cannot be properly differentiated, the arbitrary choice was made to consider 6E8S.

*Construction of the RG-2 RNA G4 initial models*

Maestro software (Maestro, Schrödinger, LLC, New York, NY, 2021.) was used to remove the third tetrad from PDB:3CDM and obtain a two-tetrad G-quadruplex, coherent with RG-2. The same software was also used to insert the two missing nucleotides in loop L1 and the missing nucleotide in loop L3. The central $K^+$ cations, necessary for the stability of G4, was also added. PyMol (The PyMOL Molecular Graphics System, Version 2.0 Schrödinger, LLC.) was used to convert the DNA structure to RNA and replace all thymine with uracil, using "Mutagenesis Wizard" tool. The hybrid RG2 model was constructed isolating the guanidine core from PDB:6E8S structure, which presents two tetrads, as our target. The loops were extracted from those obtained from the RG2 parallel G4 structure and added to the guanidine core using PyMol.

*Classical molecular dynamics protocol*

The protocol used is the same previously described.[4] The RG-2 initial model is solvated in a TIP3P[10] (80 x 80 x 80) $Å^3$ water box and charges are neutralized by adding additional $K^+$ potassium ion, using Amber *Tleap* utilities. RG-2 has been modeled with the the Amber ff99 force field[11] including the *parmbsc0-χOL3* corrections[12] to correctly represent RNA strands. The conformational space was sampled via a 1 µs molecular dynamics performed with the NAMD code,[13,14] in two replicates. A time step of 4 fs has been chosen to propagate the Newton equations of motion. This choice was possible by using the Rattle and Shake algorithm[15] combined to the hydrogen mass repartitioning scheme.[16] Langevin isotherm thermostat and piston[17,18] are used to enforce an isothermal and isobar (NPT) ensemble at a temperature of 300 K and a pressure of 1 atm. A cut off at 9 Å is used for electrostatic interaction, which are estimated using with Particle Mesh Ewald.[19] Before production MD, the system was minimized for 1000 steps, then equilibrated and thermalized by progressively removing positional harmonic constraints on heavy atoms during 36 ns. Finally, the structural parameters of the nucleic acid G4 were monitored using the script by Tsvetkov *et al.*[20] and the VMD software.[21]

*Simulation of the ECD spectra by QM/MM*

The protocol for the simulation of the ECD spectra is identical to the one used in our previous contribution.[4] 100 snapshots were randomly extracted from each MD trajectory and the eight guanines forming the two G-quadruplex tetrads, as well as the central $K^+$ ion, were included in the QM partition. The Orca software[22,23] was used to calculate the vertical transitions of each snapshot with time-dependent density functional theory (TD-DFT), using ωB97XD[24] and M06-2X[25] functionals and the 6-31G basis set. Amber software was used as an external interface to provide the QM/MM electrostatic embedding,[26] dangling bonds between MM and QM partitions have been treated with the link atom approach. The final spectrum is obtained by combining all the spectra calculated on each frame; Gaussian functions of Fixed Width at Half Maximum (FWHM) of 0.15 eV were used to convolute the spectrum using excitation energies and the rotatory strength.

### Experimental Methodology

*FRET Methodology*.

FRET experiments were performed on a 96-well format Applied Biosystems™ QuantStudio 6 PCR cycler with a FAM (6-carboxyfluorescein) filter. The sequences of RG-1 and RG-2 (5'-AGGCUGGCAAUGGCGG -3' and 5'-AGGUAUGUGGAAAGGUUAUGG -3', respectively), were modified with FAM and TAMRA (6-carboxy-tetramethylrhodamine) probes at the 5' and -3' ends (IDT, Integrated DNA Technologies). Stock solutions were prepared solubilizing the lyophilized sequences in RNAase free buffer (Merck). Stock solutions were diluted to the desired concentration using 60 mM potassium cacodylate buffer (pH 7.4). G4 folding was obtained heating the solutions to 95 ºC for 5 min, followed by slowly cooling to room temperature overnight. In the final solutions, G4s final concentration was set to 0.2 µM (total volume of 30 µl). PDS was dissolved in DMSO to give 1 mM stock solution and further diluted with the buffer reaching a total percentage of DMSO never above 0.1 %. FAM emission data were collected in duplicate in the range 25-95 ºC (with a ramp of 1 ºC every 30 s). To compare different sets of data, emission data were normalized from 0 to 1.

*Circular Dichroism*
ECD spectra were recorded on a Jasco J-715 spectropolarimeter at 25°C, adding increasing amounts of PDS to solution of RG-2 G4s at constant concentration. The parameters were the following: range 400-220 nm, response: 0.5s, accumulation: 4, speed 200 nm/min. The different buffers used are indicated in the legends of the respective figures. For this experiment we the RG-2 sequence 5'-GGUAUGUGGAAAGGUUAUGG -3'. Topology of RG-2 was identified using the conformation index r calculated as $r = CD_{265} / (|CD_{265}| + CD_{290})$ where $CD_{265}$ and $CD_{290}$ are the CD ellipticities at 265 and 290 nm, respectively. $r \geq 0.5$, $0 \leq r < 0.5$, and $r < 0$ correspond to predominantly parallel, hybrid, and antiparallel topologies, respectively.[27] The topology was also confirmed using the fitting algorithm of G-quadruplex CD spectra described by Villar-Guerra et al. implemented in the open-source R software environment.[28]

A) B)

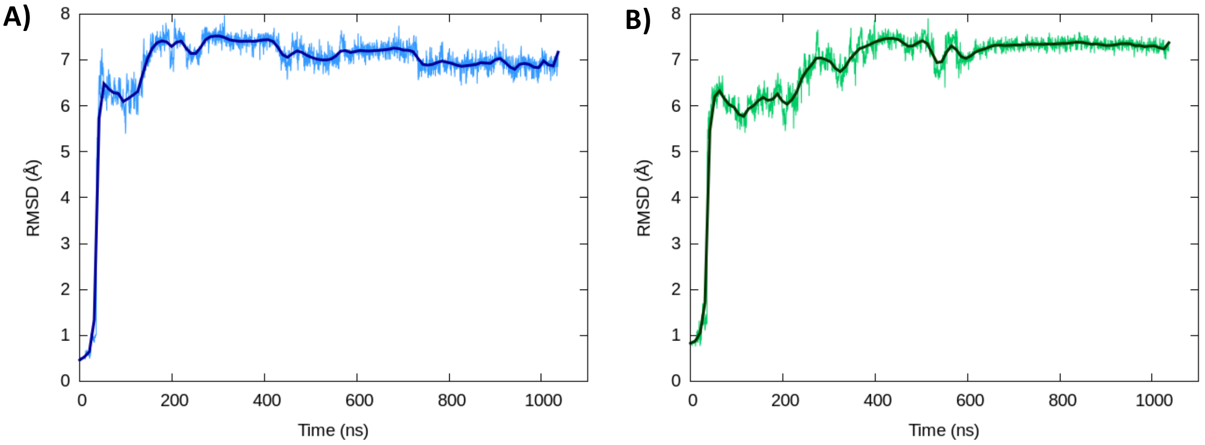


**Figure S3**. Time evolution of the RMSD of the nucleic acid for the second replica for the parallel (A) and hybrid (B) folding. The corresponding RMSD of the first replicae are given in the main text

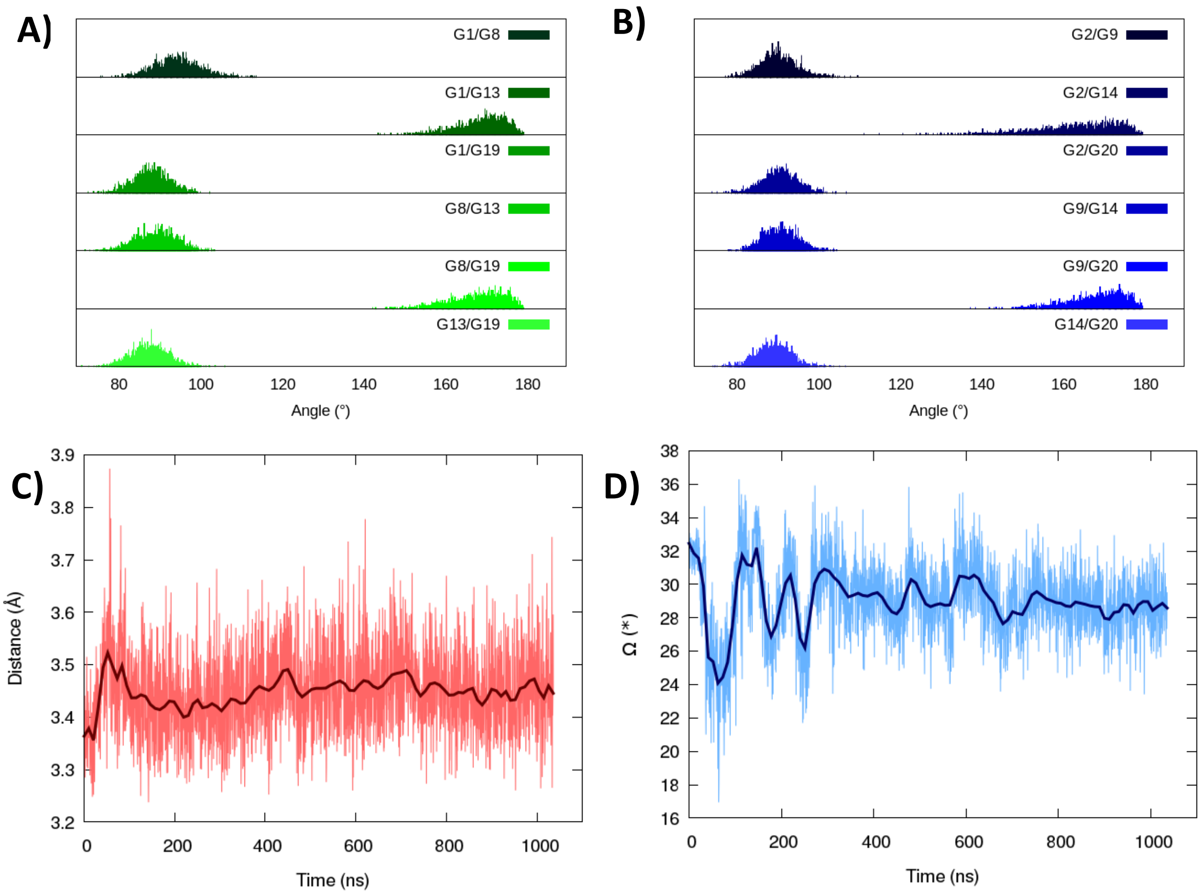


**Figure S4**. Structural parameters of the parallel RG-2 G4 obtained from the second replica. Angles between the guanine forming the first (A) and second tetrad (B), distance between the tetrads (C) and twist angle (D).

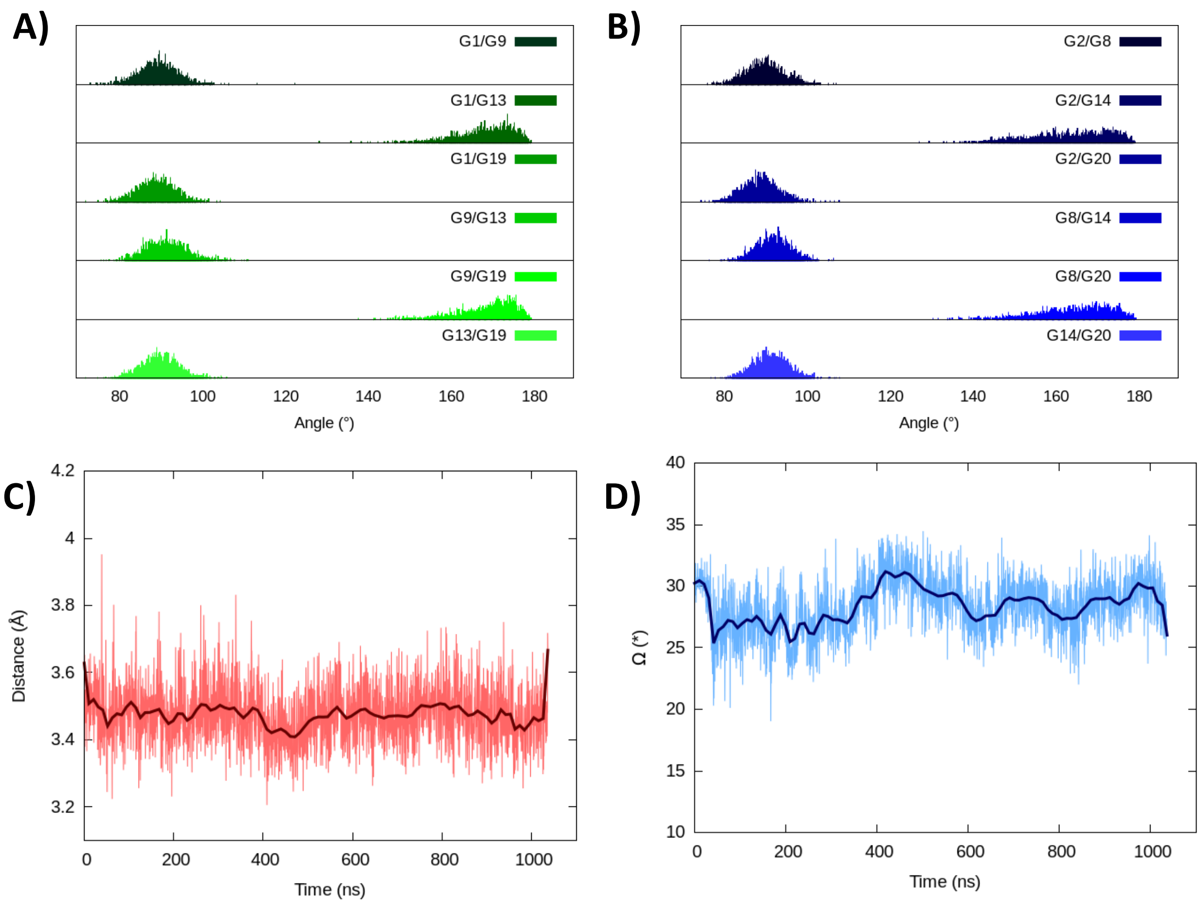


**Figure S5**. Structural parameters of the hybrid RG-2 G4 obtained from the second replica. Angles between the guanine forming the first (A) and second tetrad (B), distance between the tetrads (C) and twist angle (D).

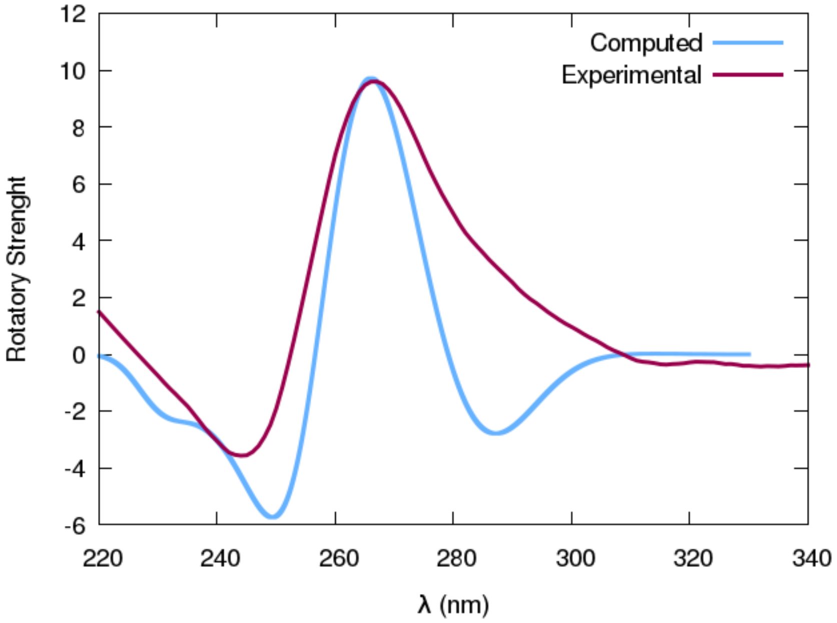


**Figure S6**. Experimental ECD spectrum of RG-2 recorded in $K^+$-containing water solution compared with the simulated spectrum obtained using the ωB97XD functional for RG-2 folded in parallel G4 topology.

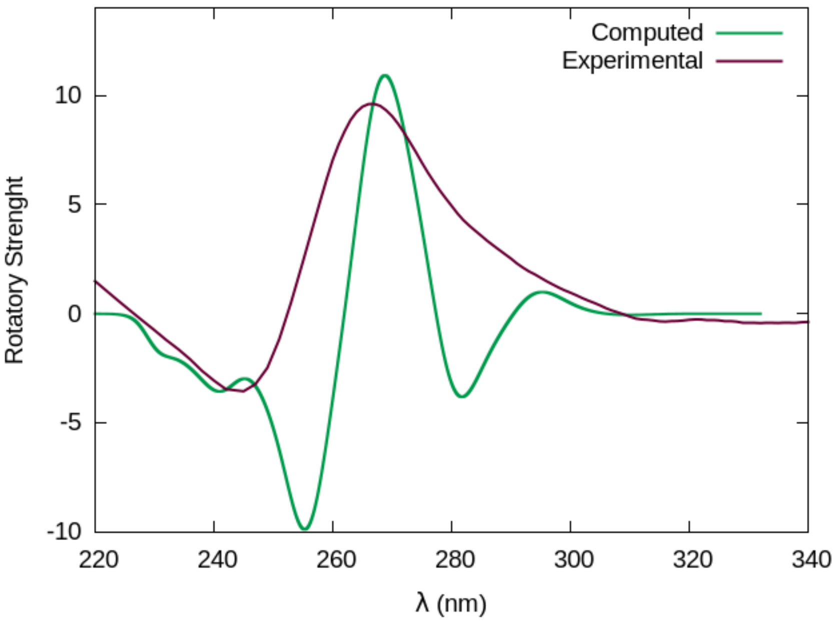


**Figure S7**. Experimental ECD spectrum of RG-2 recorded in $K^+$-containing water solution compared with the simulated spectrum obtained using the ωB97XD functional for the RG-2 folded in hybrid G4 topology.

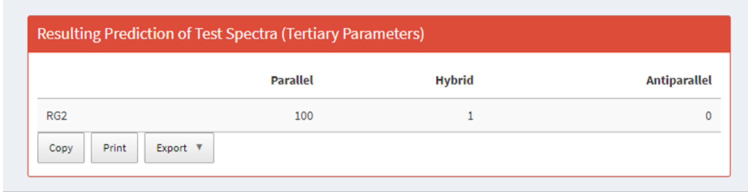

Resulting Prediction of Test Spectra (Tertiary Parameters)

| | Parallel | Hybrid | Antiparallel |
|---|---|---|---|
| RG2 | 100 | 1 | 0 |

Copy Print Export ▼

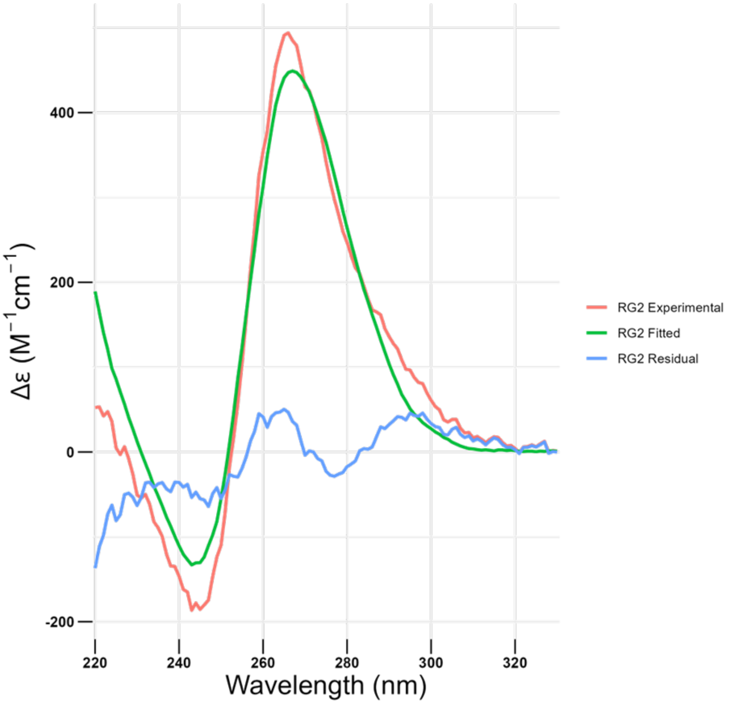


**Figure S8.** Experimental (red), fitting (green) and residual (blue) ECD spectra of RG-2 (Buffer:Tris-HCl 10 mM, KCl 150 mM). The intensity of the ECD spectra (330-220 nm) is expressed as Δε ($M^{-1}$ $cm^{-1}$) per strand.

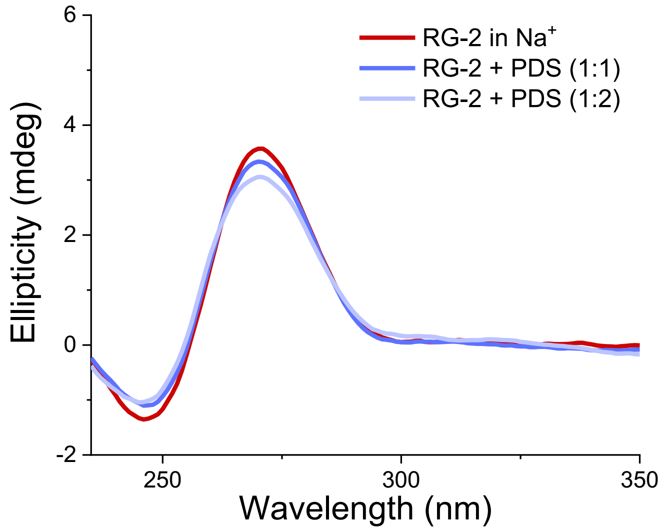


**Figure S9.** ECD spectra of RG-2 in the presence of increasing aliquots of PDS at the indicated ratios. Buffer: Tris-HCl 10 mM, NaCl 150 mM, pH=7.4

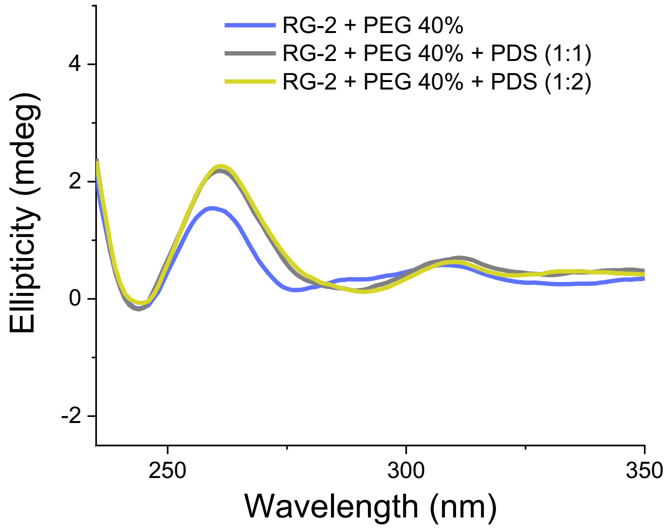


**Figure S10.** ECD spectra of RG-2 in the presence of increasing aliquots of PDS at the indicated ratios. Buffer: Tris-HCl 10 mM, KCl 150 mM, PEG 40% pH=7.4.

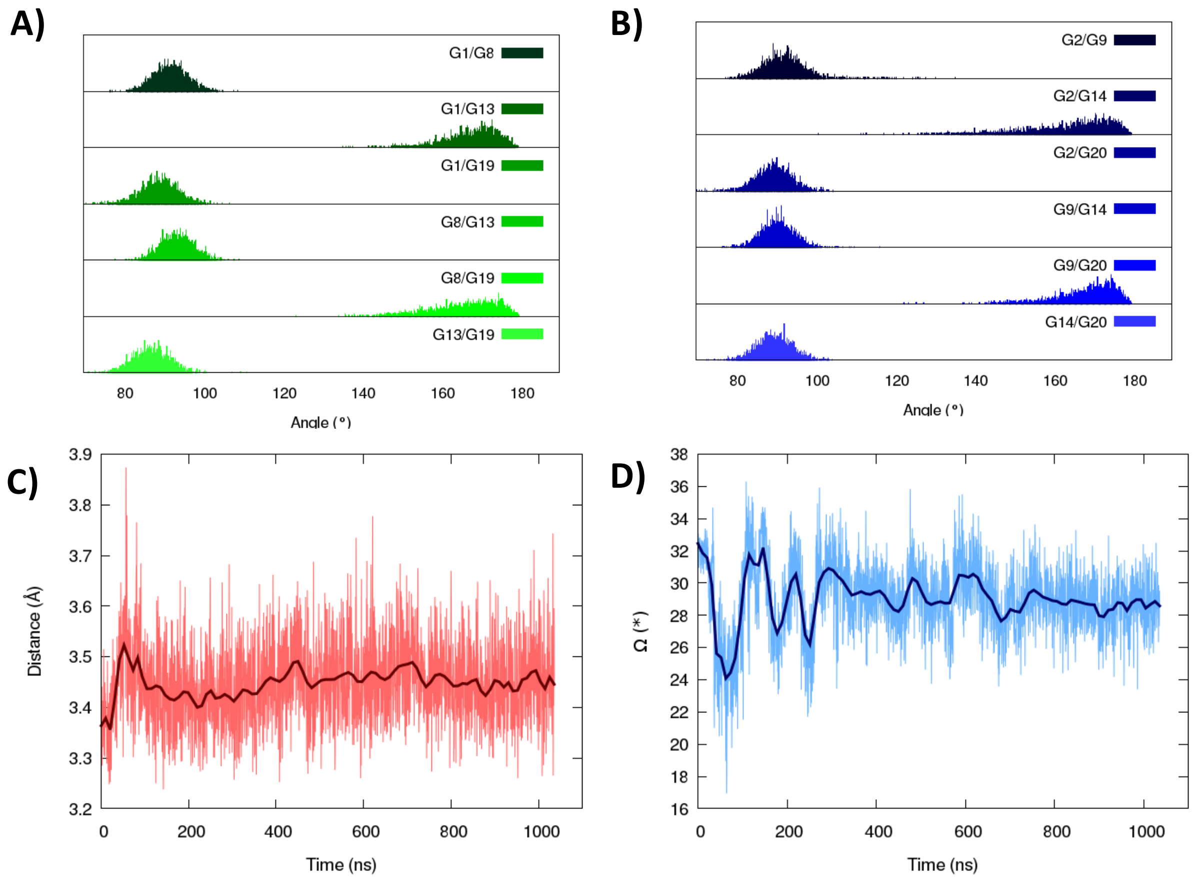


**Figure S11**. Structural parameters of the parallel RG-2 G4 in presence of PDS. Angles between the guanine forming the first (A) and second tetrad (B), distance between the tetrads (C) and twist angle (D).

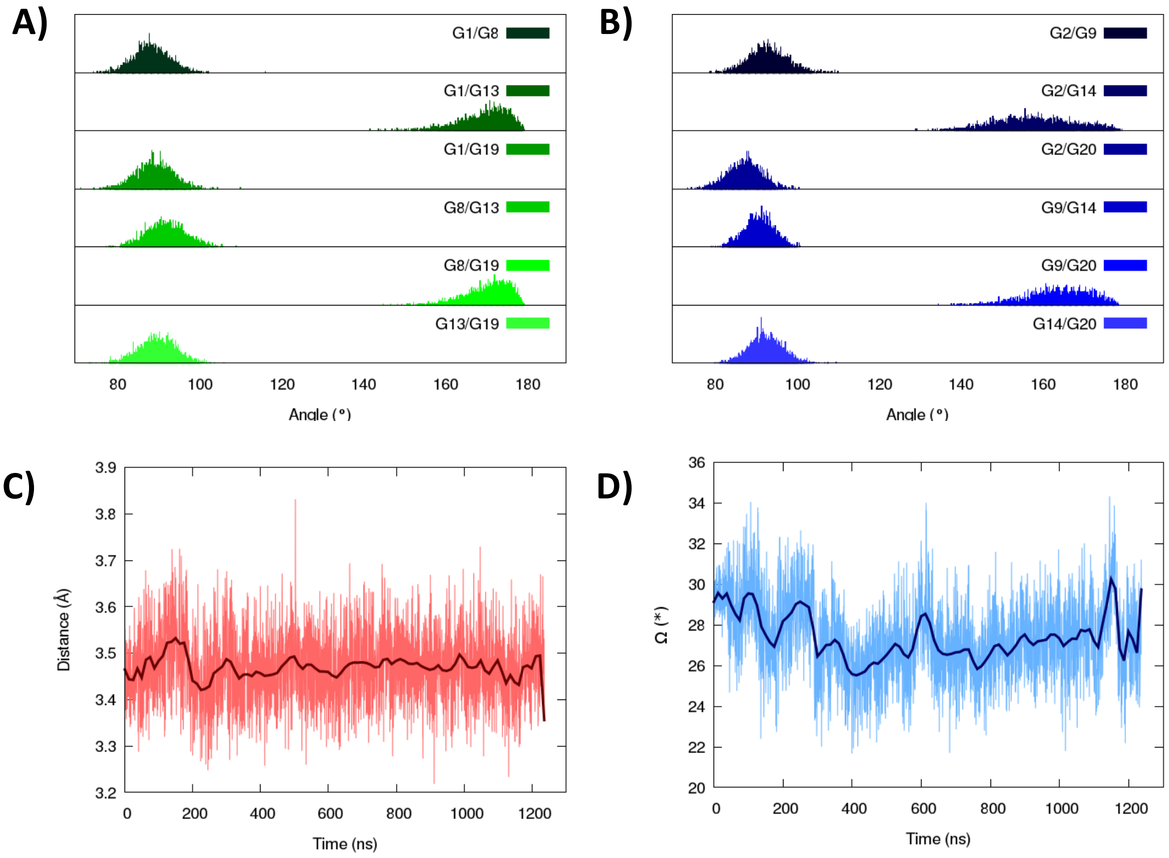


**Figure S12**. Structural parameters of the hybrid RG-2 G4 in presence of PDS ligand. Angles between the guanine forming the first (A) and second tetrad (B), distance between the tetrads (C) and twist angle (D).
